# Loss of secondary motor cortex neurons in chronic neuropathic pain

**DOI:** 10.1101/2023.01.05.522932

**Authors:** Amrita Das Gupta, Hongwei Zheng, Jennifer John, Sanjeev Kaushalya, Livia Asan, Carlo A. Beretta, Rohini Kuner, Johannes Knabbe, Thomas Kuner

**Author notes:** These authors contributed equally (Amrita Das Gupta, Hongwei Zheng).

## Abstract

Chronic neuropathic pain is associated with structural plasticity of the brain on different spatial scales, yet, little is known on the mesoscopic scale of tissue composition. Here, we determined the cellular composition of cortical areas and structural variability of entire neurons during the development of chronic neuropathic pain using longitudinal in vivo two-photon microscopy and behavioral assessment. When monitoring cell type composition in response to spared-nerve injury in 84 cortical volumes containing ∽25000 cells each, we found neuronal loss in the secondary motor cortex region M2 immediately adjacent to the cingulate cortex already one week after surgery. Loss of mostly interneurons was also evident when monitoring individual M2 neurons over time. This neuronal loss was preceded by decreased spine density and loss of distal dendritic branches. In conclusion, our work delineates M2 as a novel site and neuronal loss as a so far underappreciated mechanism underlying chronic neuropathic pain states.

## Introduction

Chronic neuropathic pain has been shown to be associated with structural alterations of the brain, ranging from macroscopic grey-matter-volume changes (Apkarian et al. 2004; Seminowicz et al. 2009; Bilbao et al. 2018; Silva and Seminowicz 2019) to microscopic alterations on the cellular level (Metz et al. 2009; Kim et al. 2011; Kim and Nabekura 2011; Kassem et al. 2013). In principle, structural neocortical changes associated with chronic pain could be based on a multitude of mechanisms. The most commonly discussed mechanism concerns structural neuronal plasticity, i.e. changes in dendritic architecture and spine density (Holtmaat and Svoboda 2009). Such changes have been shown in many cortical areas to underly functional changes in multifarious physiological and pathophysiological functions such as learning and memory, fear processing, addiction and others (Forrest et al. 2018), and can be viewed as a general disease-relevant mechanism. Changes in the microenvironment of the neuropil including extracellular space regulation (Mascio et al. 2022), ion homeostasis and others could also undergo long-lasting adaptive changes and affect neuronal communication. Such changes would influence astroglial, microglial, oligodendroglial functions and likely also endothelial cells and pericytes. Plasticity could arise from changes in structure and function of these cells, but also by changes in their distribution pattern and number. The latter is well known in neuroinflammation and neurodegeneration, where astrogliosis and proliferation of microglia is particularly well established (Salter and Stevens 2017; Ji et al. 2019). Hence, to address structural mechanisms underlying chronic neuropathic pain states, monitoring the cellular composition of cortical pain areas after induction of chronic pain is a promising avenue to follow. We recently developed an approach called “NuClear” that allows longitudinal, comprehensive, monitoring of cell type-composition in living animals (Das-Gupta et al. 2022). The resulting 3D matrix of identified cells and their location can be repetitively acquired from the identical cortical area and changes in number and location of any cell type can be identified, for example after establishing chronic neuropathic pain using the spared-nerve injury (SNI) model (Decosterd and Woolf 2000). Corresponding changes in pain behavior can be interspersed and allow correlations with structural changes.

Cortical pain processing occurs in multiple brain regions also referred to as the pain matrix (Tracey and Mantyh 2007), with each area contributing specialized functions. The cingulate cortex (CC) and its subdivisions is consistently reported to represent emotional and motor aspects of pain perception (Bliss et al. 2016; Kuner and Kuner 2021) and confers pain hypersensitivity via a direct pathway to the posterior insula (Tan et al. 2017). Studies of dendritic structural plasticity have so far addressed the somatosensory (S1) and medial prefrontal cortices (Metz et al. 2009; Kim et al. 2011; Kim and Nabekura 2011), yet has not been studied much in the CC.

Here, we attempted to address structural changes of the CC in response to SNI using NuClear and imaging of individual neurons in an in vivo longitudinal study design. In response to SNI, we surprisingly found a loss of neurons in the medial M2 cortex, immediately lateral to the CC. Most of the lost neurons were interneurons, suggesting that loss of inhibition leads to stronger activation of local pyramidal neurons. Neurons prone to die showed a loss of distal dendritic branches and a loss of dendritic spines, suggesting that changes in synaptic inputs may, together with other factors, induce neuronal cell death. In conclusion, we identified medial M2 as a novel player in chronic pain and a loss of M2 neurons as a mechanism previously not considered for chronic pain.

## Results

We assessed structural changes associated with nerve injury in the cingulate cortex (CC) and adjoining secondary motor cortex (M2) on two levels: (1) changes in tissue composition represented by the number of major brain cell types within large two-photon (2P) imaging volumes (∽0.25 mm^3^) and (2) cellular and subcellular changes on the level of entire neurons and their dendritic spines (Fig. 1A). Both levels were investigated in a longitudinal paradigm combining SNI, two-photon in vivo imaging and pain behavior (Fig. 1B). Tissue composition was determined by imaging Histon2B-GFP-labelled nuclei (Asan et al. 2021) and subsequent classification of the cell type derived from specific features of the nucleus (Das-Gupta et al. 2022). Subcellular changes were investigated by repetitively imaging individually labeled or small groups of labeled neurons in their entirety.

**Figure 1:**
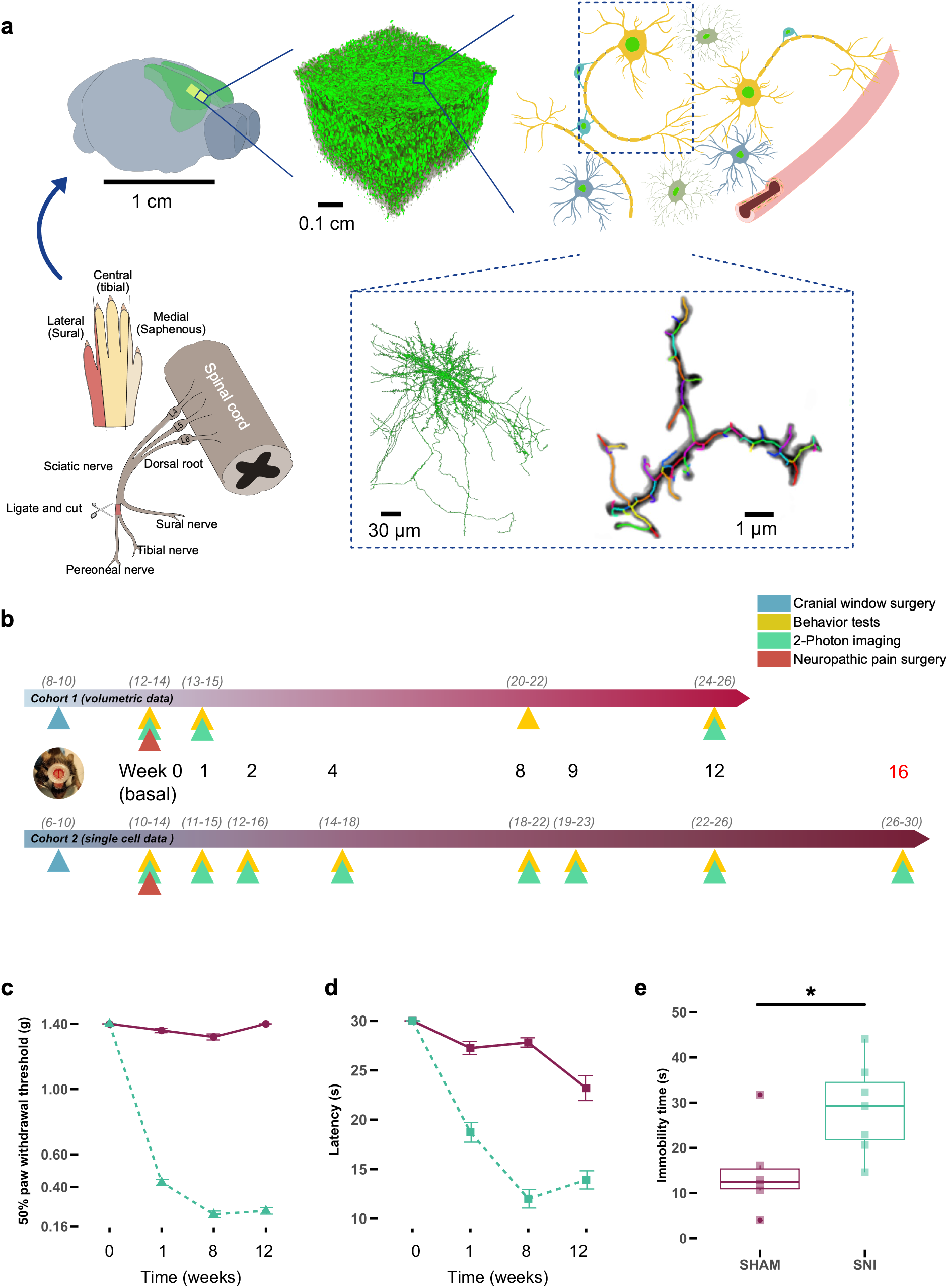
Experimental design. **(a)** A spared nerve injury (SNI) surgery was carried to induce chronic pain in mice. Effects of SNI were studied in the secondary motor area (M2) and anterior cingulate cortex (ACC) using in vivo 2-photon (2P) microscopy of cell nuclei and single neurons. Nuclei were classified into cell types using a neural network classifier. (**b**) Longitudinal study design for Cohort 1 (nuclei imaging) and cohort 2 (single neuron imaging) (**c**) von Frey test shows significant differences of the threshold filament between sham and SNI animals at 50% paw withdrawal threshold and (**d**) cold plate latency is significantly lower in SNI mice. (**e**) Immobility time showing significant difference between SNI and Sham. All behavior data N=10 for SNI and sham.

### SNI reliably induced chronic neuropathic pain states

As a foundation for the structural analysis aimed at by this study, we first report the pain behavioral readouts for both cohorts of mice studied. All mice underwent sensory withdrawal testing using von Frey filaments at the timepoints relative to the SNI or sham surgery as indicated in Figure 1B. Allodynia and hypersensitivity were detected when testing the left hind foot on which surgery was performed (Fig. 1C, Supp. Fig. 1A), but not when testing the right side (Supp. Fig. 1B). Furthermore, cold plate withdrawal was found to be significantly different in SNI-treated mice (Fig. 1D). To test the emotional aspect of pain, we assessed behavior of mice in the open field. SNI-treated mice showed longer lasting immobility, consistent with the presence of a chronic pain state (Fig. 1 E).

### Tissue composition in volumes encompassing cingulate cortex and adjacent M2 regions

After establishing the development of a chronic pain state in all mice tested, we now focus on cohort 1 mice and tissue composition analysis. We defined four 700 μm x 700 μm x 500 μm large imaging volumes in both hemispheres that were placed as close to the midline as permitted by the sagittal sinus, a structure that obstructs direct 2P imaging of brain areas residing below it (Fig. 2A, Supp. Fig. 2A). Owing to a variable position of large bridging veins towards the sagittal sinus, imaging volumes could not be placed at identical positions in each mouse (indicated by light grey squares in Fig. 2A). We took advantage of this situation by dividing the imaging volumes into cingulate/M2 and M2/M1 volumes, the latter serving as controls for the expected cingulate-specific effects. Nucleus imaging was done before, 1 week and 12 weeks after SNI surgery and yielded 20000 to 30000 cell nuclei per imaging volume (Fig. 2B). Using the NuClear approach (Das-Gupta et al. 2022), a method allowing for non-biased neuronal network-based identification of all major cell types residing within the imaging volume solely relying on a set of radiomic features of the cell nucleus (Das-Gupta et al. 2022)we determined 3D matrices of the localization of all neurons, astroglia, oligodendroglia, microglia and endothelial cells present in the volume (Fig. 2C). When applying this analysis to all 84 imaging volumes taken from 21 mice, encompassing >2 million cells, no significant changes in cell numbers were found (Fig. 2D). As expected, the most frequent cell type are neurons which were about as abundant as all other cells taken together. When re-slicing the large volumes into coronal sections, the cortical layering becomes evident and the different spatial distributions of the cell types can be appreciated. For example, neurons are organized in layers, microglia and astroglia are evenly distributed at rather constant nearest-neighbor distances (Fig. 2E).

**Figure 2:**
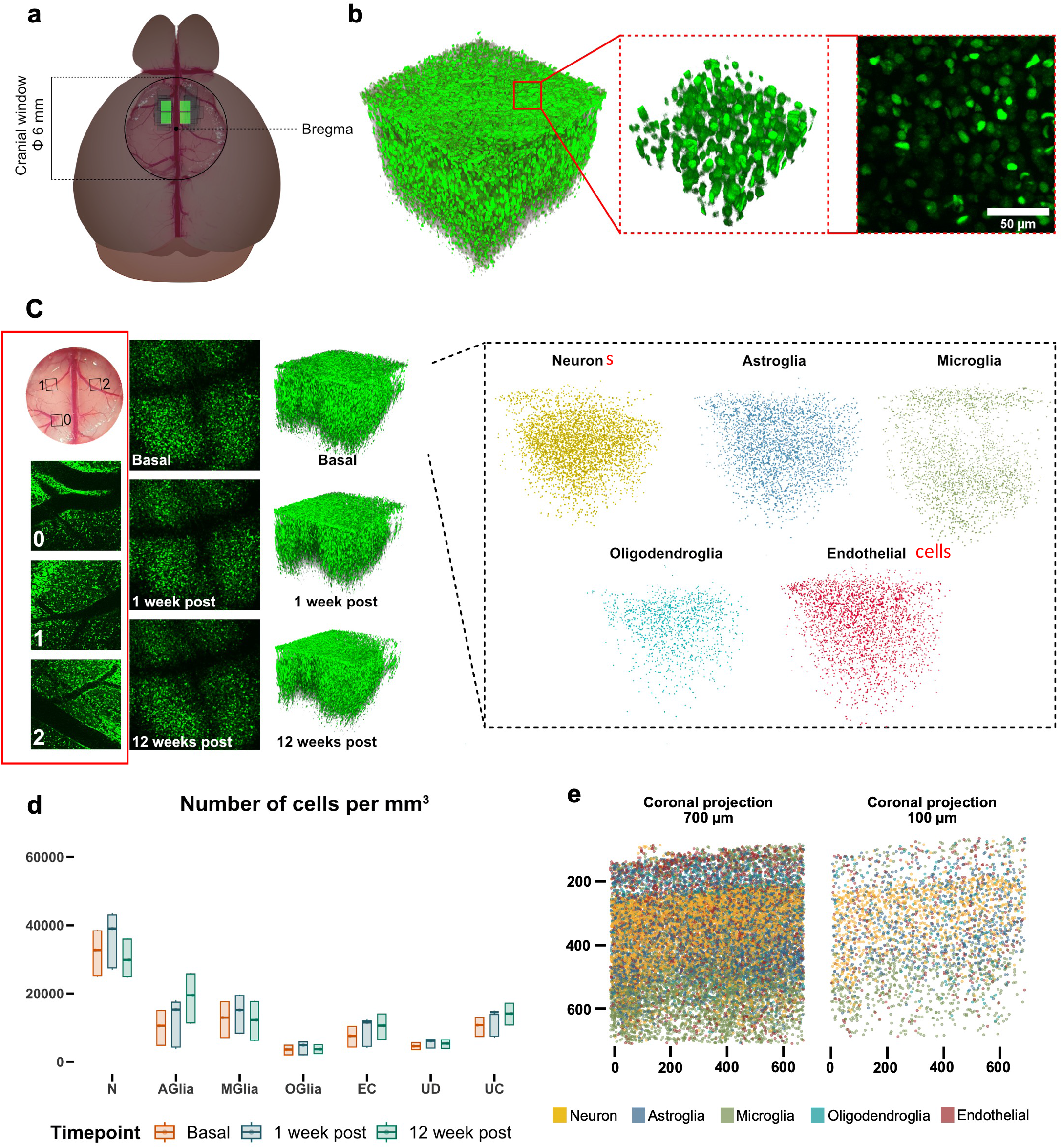
Imaging analysis of cohort 1. **(a)** Schematic representation of imaging areas for 2P imaging in cohort 1 (**b**) 3D reconstruction of a volumetric image. Zoom in shows a sub volume and a 2D maximum intensity projection (50 μm). (**c**) Imaging positions were reidentified over time using large vessels as fiducials. Each imaging volume was further classified into different cell types using a pre-trained neuronal network. (**d**) Absolute cell count per mm^3^ for different imaging timepoints. N=10 for SNI and sham. (**e**) Coronal projections (700 μm and 100 μm) showing distribution of classified cells in distinct pattens.

### Chronic neuropathic pain is associated with changes in cell number contralateral to the lesion

We next compared the number of cells in each category at different timepoints post-SNI or sham surgery. Because the medial imaging volumes contain both cingulate cortex and M2 cortex (Fig. 3, center), we digitally separated the volumes into a medial wedge representing the cingulate cortex (black) and a lateral wedge representing the adjoining M2 cortex (light blue).

**Figure 3:**
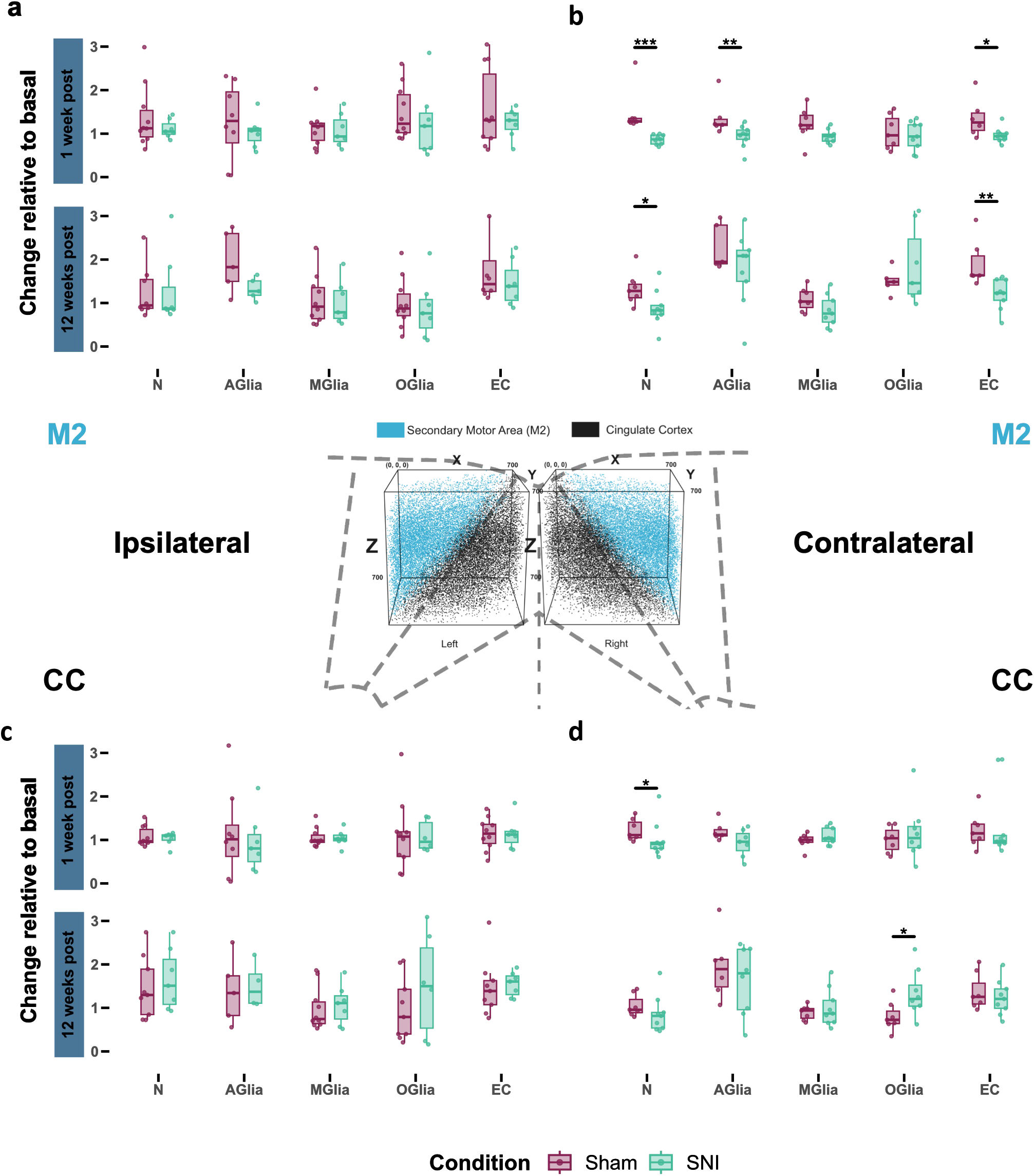
Changes in cell type composition in CC and M2. **(a)** Cell type changes in ipsilateral and contralateral hemispheres. Inlay depicts a schematic representation of the imaging volumes with regard to brain areas (dotted line), ACC nuclei are shown in blue, M2 nuclei in black. N left panel: SNI =7 / Sham = 10; right panel: SNI = 9 / Sham = 7. Wilcox test. M2: week 1 contralateral neurons: *** (p = 0.000175); Aglia: ** (p = 0.005240); EC: * (p = 0.022900); week 12: contralateral neurons: * (p = 0.03110); EC = ** (p = 0.00122). CC: Week 1 contralateral neurons: * (p = 0.0311); Week 12 contralateral Olig: * (p = 0.0418).

We identified several significant changes in cell number on the contralateral side to the lesion, where such changes are expected to occur (Fig. 3B,D). In contrast, no differences between SNI and sham were detected in ipsilateral volumes (Fig. 3A,C). Hence, we now focus on changes in cell number found for the different cell types on the contralateral side specifically affected by the lesion.

### Loss of neurons in the medial part of M2 associated with chronic neuropathic pain

A decreased number of neurons was evident already one week after SNI in the medial M2 and the CC (Fig. 3B,D). This decrease persisted throughout week 12 in M2, but not in CC. The weakly significant difference in CC may arise from an imperfect separation of CC versus medial M2 regions, with imaging volumes analysed for CC containing a certain fraction of M2 territory. The much stronger and highly significant difference in the number of neurons found in medial M2 may thereby ‘contaminate’ the CC imaging volume. Hence, our data suggests that SNI causes a persistent loss of neurons in the contralateral medial M2 cortex, and to some extent possibly also in the CC. This decrease may contribute to the chronic pain state.

### Changes in the number of non-neuronal cells

We further found a decrease in the number of astroglia in medial M2 after one week that was, however, no longer evident 12 weeks after the lesion (Fig. 3B). Microglia show a trend to be decreased in M2 as well. We further detected a significant increase in oligodendroglia in the CC after 12 weeks, while the large variability in M2 prevented a similar readout (Fig. 3B). This increase in oligodendrocyte number may suggest an activity-dependent differentiation of oligodendrocyte precursor cells (Faria et al. 2021). Lastly, we found a decrease in endothelial cell number already at week one that persisted until week 12 (Fig. 3B). This may correspond to a reduced capillary density in medial M2 and could go along with a reduced supply of oxygen and nutrients, both of which could be related to neuronal loss.

Taken together, our data introduce the medial part of M2 as a new cortical region involved in pain processing and establishes neocortical neuronal cell death as a cellular mechanism underlying chronic neuropathic pain.

### Longitudinal imaging of individually labeled neurons confirms neuronal loss in medial M2

To validate the findings obtained using the NuClear approach and to learn more about structural changes of individual neurons after SNI surgery, we sparsely labeled neurons within the contralateral medial M2 region (Supp. Fig. 3A) and repetitively imaged the same neurons over a time course of 16 weeks (see Methods). Figure 4A shows eight labeled and fully reconstructed as well as spatially registered neurons at basal state prior to SNI and at 1, 9 and 16 weeks thereafter. Evidently, only three neurons persist for 16 weeks (N6-8), but already after one week neuron 1 disappeared, followed by neurons 2, 3 and 4 at week 9 (further examples in Supp. Fig 3). On average, approximately 40% of labeled neurons were lost within 16 weeks after SNI (Fig. 4C, red curve). In sharp contrast, nearly all neurons survived over the entire imaging period in sham-treated mice (Fig. 4B, D), indicating that the loss of mGFP expression with time or toxicity triggered by high level mGFP expression over extended time periods are unlikely explanations for neuronal loss. Therefore, we conclude that neurons in the medial M2 region are lost by some mechanism of cell death triggered by SNI.

**Figure 4:**
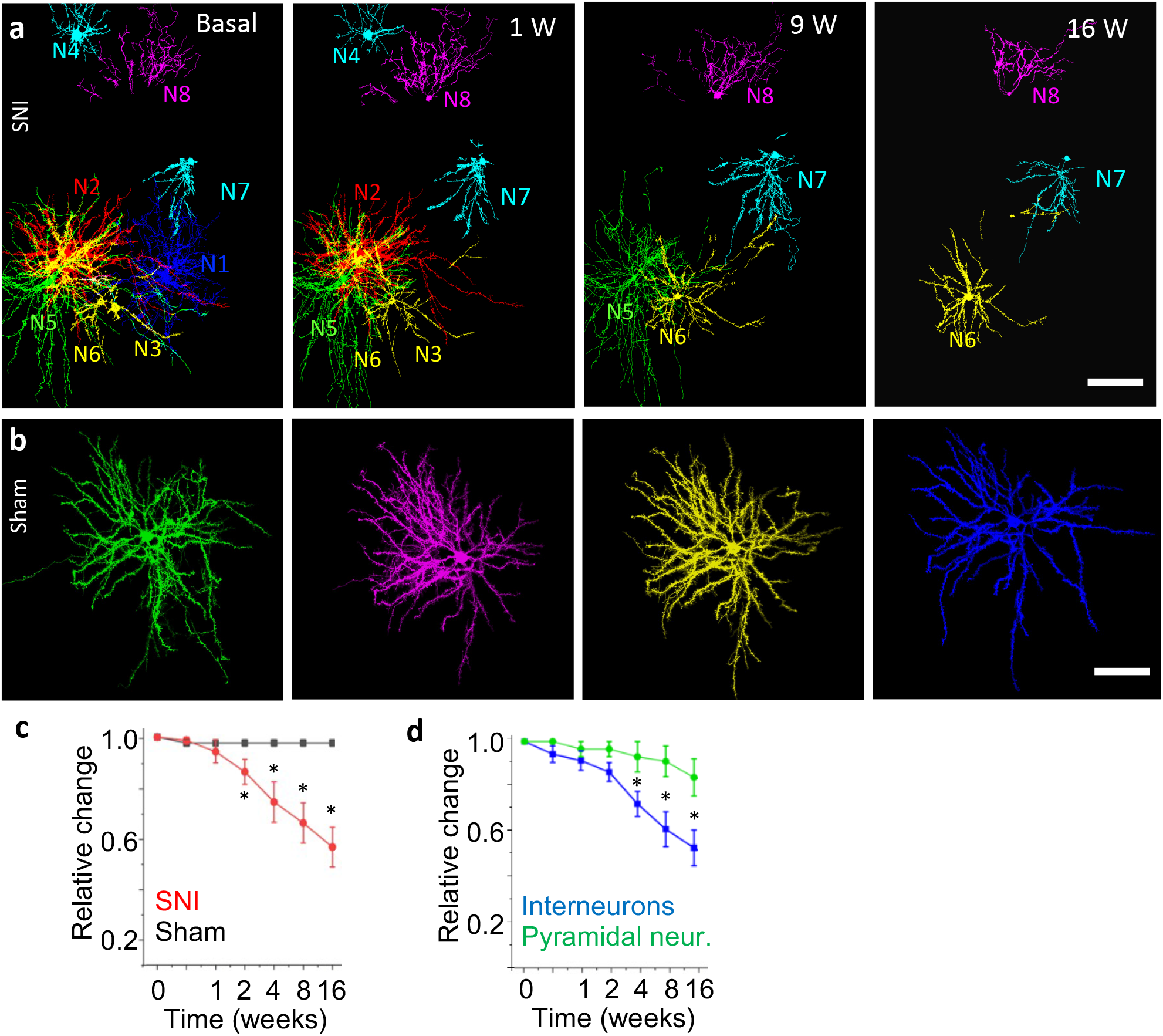
Loss of cells in medial M2 after SNI. (a) A group of eight M2 neurons imaged before SNI (basal) and 1, 9 and 12 weeks after SNI. Images show cells after image processing as summarized in Supp. Fig. 4 and the Methods supplement. Shown are fully reconstructed neurons with the raw fluorescence excised based on the traced skeleton (hence no background visible). Scale bar 100 μm. (b) Neuron imaged in M2 of a sham mouse at the same time points reported in (a). Scale bar 100 μm. (c) Quantitative comparison of the number of cells counted at different time points in sham and SNI treated mice. N=20 cells for sham and N=28 for SNI, taken from a total of 16 mice. Mean ± SEM. ANOVA with Bonferoni multiple comparison test, * depict P <0.05. (d) Quantitative comparison of the number of lost interneurons versus pyramidal neurons after SNI. N=14 pyramidal cells and N=18 interneurons, taken from a total of 16 mice. Mean ± SEM. ANOVA with Bonferoni multiple comparison test, * depict P <0.05.

### Interneurons are lost more frequently than pyramidal neurons

A differential loss of interneurons versus pyramidal neurons could have a strong impact on network function, with a loss of interneurons reducing local inhibition and thus making pyramidal cells more active, and a loss of pyramidal neurons attenuating the output of the neuronal ensemble in the medial M2. To address this, we sorted the imaged neurons according to morphological parameters into interneurons and pyramidal neurons (see Methods). Our analysis revealed that interneurons are more prone for cell death than pyramidal neurons (Fig. 4D). Hence, a more pronounced loss of interneurons predicts a reduction in inhibitory drive within the local M2 network.

### Structural changes of the dendritic tree precede neuronal loss after SNI

To assess if neurons prone to die undergo some prodromal state of structural changes, we analysed their entire dendritic tree including their postsynaptic spines (see Supp. Fig. 4 for illustration and Methods Supplement for detailed description). For example, neuron 4 (already displayed in Figure 4) underwent a pronounced loss in spine density already within the first week after SNI and barely retained any spines one week later (Figure 5A), to finally disappear before week four. In other neurons we found rearrangements of the dendritic tree. Figure 5B shows neurons 2 and 5 (as in Fig. 4). Each panel shows both neurons at the earlier time point in red and at the later time point in green, hence, structures appearing in yellow exist at both time points while red alone indicates loss of a structure. The pink rectangle marks a dynamic region while the blue rectangle delineates a rather stable region. While dendritic structures are mostly overlapping between basal and day 3 (Fig. 5B1), a loss of distal dendritic branches is evident at week 1 (Fig. 5B2) and progresses further in the delineated area (Fig. 5B3,4). At week 7, neuron 2 has disappeared (Fig. 5B5,6). Similar observations were made in all neurons analyzed (Supp. Fig. 5). In conclusion, before disappearing, neurons undergo structural changes ranging from a loss of spines to a loss of dendritic branches. These structural changes may reflect alterations in synaptic inputs, possibly triggered by SNI.

**Figure 5:**
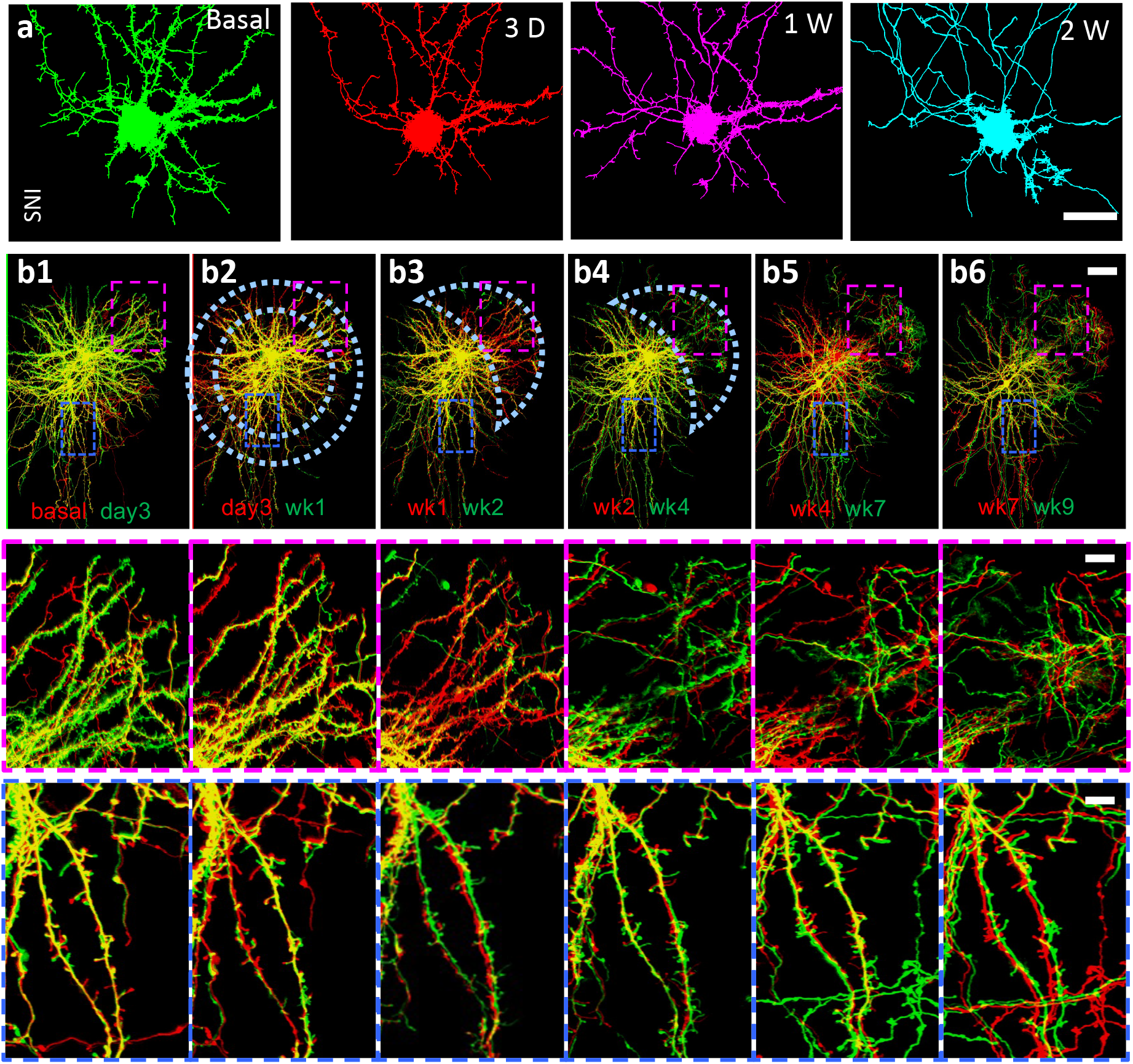
Spine loss and changes in dendritic morphology. (a) Neuron 4 (from Fig. 4) imaged at basal state, 3 days, 1 week and 2 weeks after SNI. Neuron disappeared before week 4. Scale bar 20 μm. (b) Neurons 2 and 5 (from Fig. 4) imaged at multiple time points as indicated. The respectively earlier time point is shown in red, the later time point in green, so that preserved cellular morphology will display yellow, but lost structures appear in red. The two color-coded insets (scale bars 20 μm) show details of a dynamic region (magenta) and a static region (blue). Scale bar in upper panels = 50 μm.

## Discussion

Using NuClear, an unbiased and comprehensive tissue analysis approach, we monitored the cellular composition of the cingulate and M2 cortices after SNI. While monitoring a large tissue volume (in total approximately 21 mm^3^) distributed over the medial part of both hemispheres, we found significant changes in cellular composition solely within a small subvolume encompassing the medial part of M2. The most prominent change was a reduction of the number of neurons in medial M2, but also changes in non-neuronal cells, in particular endothelial cells, were observed. The loss of neurons could be validated by longitudinally imaging individually labeled neurons in medial M2. Up to 40% of labelled neurons disappeared, mostly affecting interneurons. Neurons prone to die lost dendritic branches and spines beforehand. In conclusion, our work adds the medial M2 cortex as a novel player in chronic pain processing and suggests selective loss of cortical neurons as a mechanism of chronic pain.

### What is particular about the medial M2?

The medial M2 in rodents is considered to belong to the mPFC and is thought to mediate context-dependent and choice-related decisions, to map sensory cues to motor actions, and to potentially also contribute to depressive-like behavior (Barthas and Kwan 2017). Consistent with these diverse functions, this area receives abundant multimodal sensory inputs and projects to sensory, parietal and retrosplenial cortices (Barthas and Kwan 2017). This constellation of functionalities and in particular connectivity makes a role of the medial M2 in pain processing highly plausible, consistent with reports on patients suffering from complex regional pain syndrome (Maihöfner et al. 2007; Jung et al. 2019) However, the detailed connectivity of cells in medial M2 lost after SNI, both in terms of afferent and efferent connections, changes in activity in these projections and their impact on pain behavior await further elucidation.

### A neurodegenerative mechanisms contributing to chronic pain?

Our finding of neuronal loss starting as early as one week after SNI suggests that a mechanism of neuronal cell death may get triggered by SNI-induced changes in activity of neurons upstream of M2. The distinct reduction in spine density and loss of distal dendritic branches observed within one week after SNI further supports the idea that the loss of certain synaptic inputs or their altered activity patterns are involved in early steps, thereby eventually resulting in a denervation of distinct M2 neurons. Such loss of afferent input has been reported to result in cell loss (Mostafapour et al. 2000; Inquimbert et al. 2018). Persistent changes in activity, either silence or overactivity, are known to trigger different mechanisms of cell death (Chi et al. 2018; Fricker et al. 2018; Moujalled et al. 2021). Which of these are active in SNI-induced loss of M2 neurons remains to be investigated. Alternatively, or synergistically, the neuronal microenvironment defined by glia and endothelial cells might respond to the SNI impact and affect neuron survival. A decreased number of endothelial cells may reflect changes in microvasculature that in turn may negatively affect neuronal fidelity. In conclusion, our work implies neurodegeneration as a mechanism underlying chronic pain states, thereby extending a concept so far limited to the peripheral nervous system (Reichling and Levine 2011) to neocortical circuits.

### How can neuronal loss contribute to the development of a chronic pain state?

If neurons get lost, they can no longer contribute to function, yet chronic pain is an active percept generated by as yet unknown interactions of brain areas. As a simple approximation, an increased firing of excitatory M2 output neurons could elicit activity in other areas of the pain matrix and thereby contribute to the pain percept. Likewise, disinhibition via long-range inhibitory connections could achieve the same. Given that we found a more pronounced loss of interneurons within M2, the surviving pyramidal neurons may become more active and thereby contribute to eliciting a pain percept. Interestingly, such local disinhibition in the anterior cingulate network has been described after nerve injury (Blom et al. 2014). Likewise, hyperactive pyramidal neurons in the anterior cingulate have been linked to neuropathic pain (Zhao et al. 2018). In conclusion, while published work supports the mechanisms discussed here, additional work will be needed for a deeper understanding.

### Role of non-neuronal cells

In addition to changes in neuronal number we found a consistent change in endothelial cell number. Limited to the medial M2 region, the number of endothelial cells was reduced in SNI-treated mice compared with sham. This effect was already evident at week one and persisted until week 12. At the moment, we can only speculate about the pathophysiological contribution of this change. A possible explanation could be that the microvascular network becomes more sparse after SNI. This could in turn affect the metabolic support of M2 neurons and may be a factor synergistically acting with the structural changes in dendrites and concomitant connectivity. Astroglia were significantly reduced after SNI at week one, but not at week 12. This could be an apparent difference related to the higher variability of the data at week 12. However, the lack of astrogliosis would rather argue against a neurodegenerative mechanism of neuronal cell death in M2 after SNI.

### Technical aspects of our imaging and analysis approach

The approach used for comprehensive tissue analysis, NuClear, delivers an unbiased and comprehensive analysis of cell type composition. The number of cell types detected depends on the training data used for the neuronal network and is not limited to the cell types chosen here. The detection accuracy differs between cell types and ranges from 95% for glial cells to 99% for neurons and endothelial cells. By precisely re-identifying imaging volumes acquired at different time points, artificial alterations of cell numbers by misregistration is unlikely to happen. Variations in imaging quality between individual session may in some cases affect the outcome. However, given the large number of volumes analyzed here, such an effect would cancel out given the study design used here (SNI versus control, volumes ipsi- and contralateral to the lesion, medial versus more lateral volumes).

## Materials and Methods

### Ethical approval

The entire study was conducted in line with the European Communities Council Directive (86/609/ECC) to reduce animal pain and/or discomfort. All experiments followed the German animal welfare guidelines specified in the TierSchG. The study has been approved by the local animal care and use council (Regierungspräsidium Karlsruhe, Baden-Wuerttemberg) under the reference numbers G206/11, G294/15 and G27/20. All experiments followed the 3R principle and complied with the ARRIVE guidelines (Percie du Sert, Hurst et al. 2020).

### Animals

Transgenic mice expressing the human histone 2B protein (HIST1H2B) fused with enhanced green fluorescence protein (eGFP) under the control of the chicken beta actin promoter (CAG-H2B-eGFP) (Hadjantonakis and Papaioannou 2004) (B6.Cg-Tg(HIST1H2BB/EGFP)1Pa/J, Jackson Laboratory; # 006069) were used in animal experiments of cohort 1 (21 mice in total). C57BL6-J mice were used for cohort 2 (16 mice in total).

### mGFP expression in single neurons (cohort 2)

Two adeno-associated virus constructs were used: (1) A DIO cassette containing the synapsin promotor and membrane-bound GFP to achieve long-term neuron-specific expression, (2) a cassette expressing Cre-recombinase under the control of the CBA promotor. The in house-produced AAV1/2 viral particles were stereotaxically injected into the medial M2 region before the cranial window surgery using a high titer of construct (1) and a 10000-20000x dilution of construct (2), both virus solutions were mixed at equal amounts. Stereotaxic coordinates were AP: 1.8 or −0.1 to −0.5, ML: −0.25 to −0.3, DV: – 0.35. This strategy resulted in sufficiently strong expression of mGFP in a few neurons within the target volume.

### Chronic cranial window implantation

A craniectomy procedure was performed on each mouse to enable in vivo two-photon imaging following a predefined protocol as described before (Holtmaat, Bonhoeffer et al. 2009). Briefly, the mice were anesthetized with an intraperitoneal injection (i.p.) of a combination of 60 μl Medetomidine (1mg/ml), 160 μl Midazolam (5 mg/ml) and 40 μl Fentanyl (0.05 mg/ml) at a dosage of 3 μl/g body weight. The head was shaved, and the mice were fixed with ear bars in the stereotactic apparatus and eye ointment was applied. Xylocain® 1 % (100 μl, Lidocaine hydrochloride) was applied under the cranial skin and 250 μl Carprofen (0.5 mg/ml) was injected subcutaneously (s.c.). The skin was removed to expose the skull and the surface of the skull was made rough to allow the cement to adhere better. A skull island (approx. 6mm ∅) was drilled centered at bregma using a dental drill and removed with a forceps (#2 FST by Dumont) making sure not to damage the cortical surface. For improved imaging condition, the dura was carefully removed using a fine forceps (#5 FST by Dumont). Normal rat ringer solution (NRR) was applied on the exposed brain surface to keep it moist, and a curved cover glass was placed on top to cover it (Asan, Falfan-Melgoza et al. 2021). With dental acrylic cement (powder: Paladur, Kulzer; activator: Cyano FAST, Hager & Werken GmbH) the cover glass was sealed, and excess cement was used to cover the exposed skull and edges of the skin. A custom designed holder was placed on top of the window and any gaps were filled with cement to ensure maximum adhesion with the skull. After the procedure, the mice were injected (i.p. / s.c.) with a mix of 30 μl Atipamezole (5 mg/ml), 30 μl Flumazenil (0.1 mg/m) and 180 μl Naloxon (0.4 mg/ml) at a dosage of 6 μl per gram body weight. To ensure proper recovery of the mice, 3 more doses of Carprofen were given every 8-12 hours and the mice were placed onto a heating plate and monitored.

### In vivo two photon imaging

Two photon imaging was carried out with a two-photon microscope (TriM Scope II, LaVision BioTec GmbH) equipped with a tunable pulsed Titanium-Sapphire (Ti:Sa) laser (Chameleon Ultra 2; Coherent). Prior to each imaging session, the laser power was adjusted to achieve the best signal to noise ratio. Adaptations were made to minimize the effect of laser attenuation due to tissue depth by creating a z-profile and setting the laser power for different imaging depths while making sure to minimize oversaturation of image pixels.

### In vivo two photon imaging of the H2B-GFP mice (cohort 1)

Imaging of H2B-GFP was performed at a laser wavelength of 960 nm. A water immersion objective (16x; NA 0.8, Nikon) was used to obtain volumetric 3D stacks of the H2B-GFP mice. Individual frames consisted of 694 μm x 694 μm in XY with a resolution of 0.29 μm/pixel. Stacks were obtained at varying depths up to 702 μm from the cortical surface with a step size of 2 μm in Z. Mice were initially anesthetized using a vaporizer with 6% isoflurane and eye ointment was applied. For imaging, mice were fixed in a custom-built holder on the microscope stage. Isoflurane (Baxter) was adjusted between 0.5 – 1.5% depending on the breathing rate of each mouse to achieve a stable breathing rate of 55 – 65 breaths per minute with an oxygen flow rate of 0.9 – 1.1 l/min. A heating pad was placed underneath the mouse to regulate body temperature. An infrared camera was used for monitoring of the mice during the imaging.

### In vivo two photon imaging of single cell labelled mice (cohort 2)

Imaging of membrane-bound GFP (mGFP) was performed at a laser wavelength of 960 nm. A water immersion objective (25x, NA 1.1, Nikon) was used to obtain overlapping volumetric 3D stacks of whole neurons labelled with mGFP. Individual frames covered an XY area of 350 μm x 350 μm, corresponding to a resolution of 85 nm/pixel when sampling at 4092 × 4092 pixels at 16-bit gray scale resolution. Using a Z-step size of 1 μm, image stacks consisted typically of 500 sections. Four to six adjacent image stacks were acquired with a 15-20% overlap for stitching. For stitching, the resolution was reduced by 50%, typically yielding superstacks of approximately 3800 × 5900 × 500 pixels. Further details are provided in Supplementary Methods.

Mice were anesthetized using an intraperitoneal injection (i.p.) of a combination of 60 μl Medetomidine (1mg/ml), 160 μl Midazolam (5 mg/ml) and 40 μl Fentanyl (0.05 mg/ml) at a dosage of 3 μl/g body weight. For imaging, mice were fixed in a custom-built holder on the microscope stage. After the imaging session, mice were injected (i.p. / s.c.) with a mix of 30 μl Atipamezole (5 mg/ml), 30 μl Flumazenil (0.1 mg/m) and 180 μl Naloxon (0.4 mg/ml) at a dosage of 6 μl per gram body weight. For recovery, mice were kept on a heating plate.

### SNI surgery

For the induction of neuropathic pain, a spared nerve injury (SNI) of the sciatic nerve was performed after baseline imaging as described elsewhere (Gangadharan et al. 2022). Briefly, mice were anesthetized using isoflurane as described above. After applying a local anesthetic, an incision on the left hindpaw was made above the sciatic nerve. The peroneal and tibial nerve were ligated and cut and the sural nerve (innervating the lateral part of the ipsilateral paw) was left intact.

### Behavior analysis

For habituation, mice were placed in the dark behavior room for 3 days, one hour each prior to the day of the experiment as well as the day of the experiment. All behavior experiments were performed in the dark cycle of the circadian rhythm of the mice under red light conditions.

### von Frey test

To test hypersensitivity after chronic pain induction, the von Frey test was used as described elsewhere (Gangadharan et al. 2022). Briefly, all mice were habituated on the von Frey grid prior to the experiment. Each mouse was stimulated on the lateral part of both the ipsilateral and contralateral paw with von Frey filaments ranging from 0.07 g to 1.4 g. Each filament test consisted of 5 trials and trials were either marked 0 (negative response) or 1 (positive response) depending on whether the mouse showed any response like paw flicking, licking or flinching behavior. For analysis, a mouse was classified as being responsive to a certain filament when more than 50% of the trials were positive.

### Cold plate test

To test allodynia, the cold plate test way used (Tan et al. 2017). The cold plate was set at 4°C and mice were lowered onto the plate and the latency for a reaction was tested for 30s. The time for a first aversive response (paw withdrawal, jumping, licking) was noted and any mouse not responding during the first 30 s was assigned a latency of 30 s.

### Open field test

The open field test was used to test for the emotional aspect of pain. The setup consists of a 50cm x 50cm box with 15 cm high walls and a separate distinction for the center of the box. Mice were placed in the center of the box and were subsequently recorded with a video camera for 5 minutes. A custom script was designed in Bonsai for mouse tracking and analysis of time spent in the center or outside the center of the box. Immobility time excluding grooming was manually scored by an investigator blinded for the mice.

### Reidentification of stack positions over time in cohort 1

To be able to compare cell type changes in the same volume over time, the imaging positions of each imaging volume had to be identical. To achieve this, three or more large vessels on the cortical surface were used as fiducials. The center of the first vessel was set as zero on both X and Y coordinates of the microscope stage. The coordinates of the subsequent vessels and stacks were noted with respect to the first vessel. For imaging of the subsequent timepoint, the coordinates were used to identify the approximate location of the stack. After roughly reidentifying the imaging position, the microscopi field of view in different imaging planes along the Z-axis was aligned to fit to the previous position. A registration system based on the distance and angle of the first vessel from bregma was created using a self-written script in Python, which was used to assign the position of all imaging volumes with respect to bregma. Using a mouse brain atlas (Paxinos & Franklin, 2012), imaging volumes were selected which were located in the secondary motor area (M2) and the cingulate cortex (CC). Each imaging volume was divided into diagonal wedges (Fig. 3) corresponding the M2 and CC regions for both ipsilateral and contralateral hemispheres for further analysis. Due to a “pseudo”-dura consisting of probably fibroblasts growing on top of the brain, imaging was started from ∽76 μm below occurrence of the first fluorescent signal. Imaging was performed until 550 μm below the first fluorescent signal due signal-to-noise ratio increasing with depth.

### Image analysis of individual neurons used in cohort 2

The image analysis pipeline is illustrated in Supp. Figure 4 and described in detail in the Methods supplement.

Morphometric sorting of neurons: Interneurons were identified based on their multipoloar shape, lower spine density and small size. Pyramidal neurons were identified by the typical pyramidal shape of the soma and a strong apical dendrite, high spine density, and large size.

### Antibody staining

Antibody staining was performed on 50 μm thick brain slices with following antibodies: Anti-NeuN (monoclonal mouse) antibody (Synaptic Systems; #266 011; dilution 1:500), Anti-IBA1 (polyclonal rabbit) antibody (Fujifilm – Cellular dynamics; #019-19741; dilution 1:250), Anti-S100ß (polyclonal chicken) antibody (Synaptic Systems; #287 006; dilution 1:500) for astroglia. Secondary antibodies were used with 1:500 dilution.

### NuClear analysis for identification of cell types belonging to individual H2B nuclei

Details of this method are published elsewhere (Das-Gupta et al. 2022). Briefly, automated segmentation of all nuclei was performed with the StarDist neuronal network using a high performance cluster environment (bwForCluster MLS&WISO). Afterwards, radiomics features for every segmented nucleus were extracted using the PyRadiomics library and self-written python scripts. Every segmented nucleus was classified with five pre-trained neuronal network classifiers and assigned a unique class (neurons, astroglia, microglia, oligodendroglia, endothelial cells). Nuclei that were assigned to two or more classes were assigned to the “undecided” class; nuclei that were assigned to no class were labelled as “unclassified”.

### Statistical analysis

In general, statistical analysis was performed in R (R. Core Team 2021). Depending on the distribution of the data either t-tests or non-parametric statistical test (Wilcoxon signed-rank test) were used for statistical testing, p-values were corrected for multiple comparisons using the “Bonferroni”-method. Plots were created using the R package “ggplot2” (Wickham 2016).

### Statistical analysis of 3D cell type distribution

All data were stored in a local MySQL database (MySQL Workbench 8.0). Data were imported into R. Cell count and first nearest neighbor distance were calculated for each cell type for different timepoints. The nearest neighbor distance was calculated using the “spatstat” package for R (Baddeley and Turner 2005).

## Contributions

ADG: Cohort 1. Cranial window and SNI surgeries, imaging data acquisition, implementation and development of analysis pipeline, data analysis, conceptual inputs, generation of figures. HZ: Cohort 2. Analysis of 2P in vivo imaging data, development of automated tracing algorithm compatible with large data volumes, generation of figures.

JJ: Cohort 1. Cranial window and SNI surgeries, behavior and imaging data acquisition, tissue volume reidentification.

LA: Cohort 1. Development of the initial analysis pipeline for nucleus-based cell type identification, tissue volume reidentification script.

CAB: Cohort 1. Development of segmentation, application of StarDist.

SK: Cohort 2: Cranial window implantations, SNI surgeries and 2P imaging data acquisition. RK: Initial conceptual input.

JK: Conceptualization, implementation, experimentation (Cohort 1 and Cohort 2), analysis, writing of the MS, project management and supervision.

TK: Project idea and concept, writing of the MS, supervision.

## Acknowledgements

We want to thank Michaela Kaiser, Dunja Baumgartl-Ahlert and Nadine Gehrig for their excellent technical support during preparation and execution of experiments. We thank Deepitha Männich for help with posthoc analysis. We gratefully acknowledge the support by the German Research Foundation (DFG) (SFB1158, project B08 awarded to TK), the data storage service SDS@hd, supported by the Ministry of Science, Research and the Arts Baden-Württemberg (MWK) and the DFG through grant INST 35/1314-1 FUGG, as well as the high performance cluster bwForCluster MLS&WISO and HELIX, supported by the MWK and the DFG through Grant INST 35/1134-1 FUGG.

## Figure legends

**Supplementary Figure 1:**
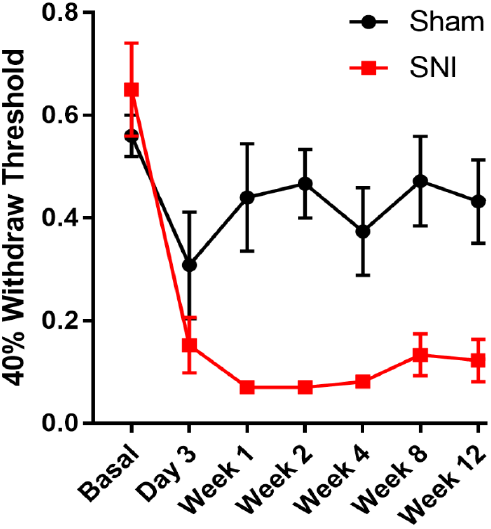
Sensory testing of cohort 2 mice. Von Frey withdrawal thresholds for SNI (N=7) and sham mice (N=6). Mean ± SEM.

**Supplementary Figure 2:**
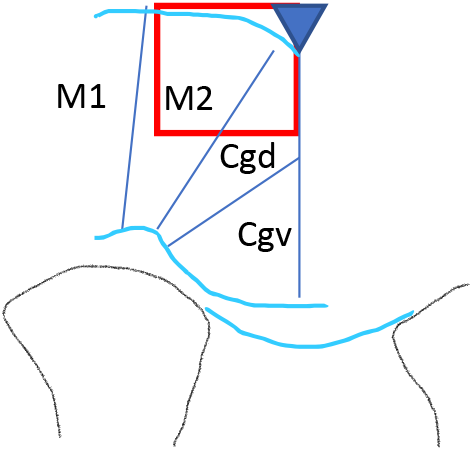
Scheme illustrating the imaging situation at the cingulate cortex. The red square represents the imaging volume of 700 μm width. The triangle demarks the sagittal sinus. Drawing approximately to scale. AP position of coronal section at approximately −0.2. M1 primary motor cortex, M2 secondary motor cortex, Cgd dorsal cingulate cortex, Cgv ventral cingulate cortex.

**Supplementary Figure 3:**
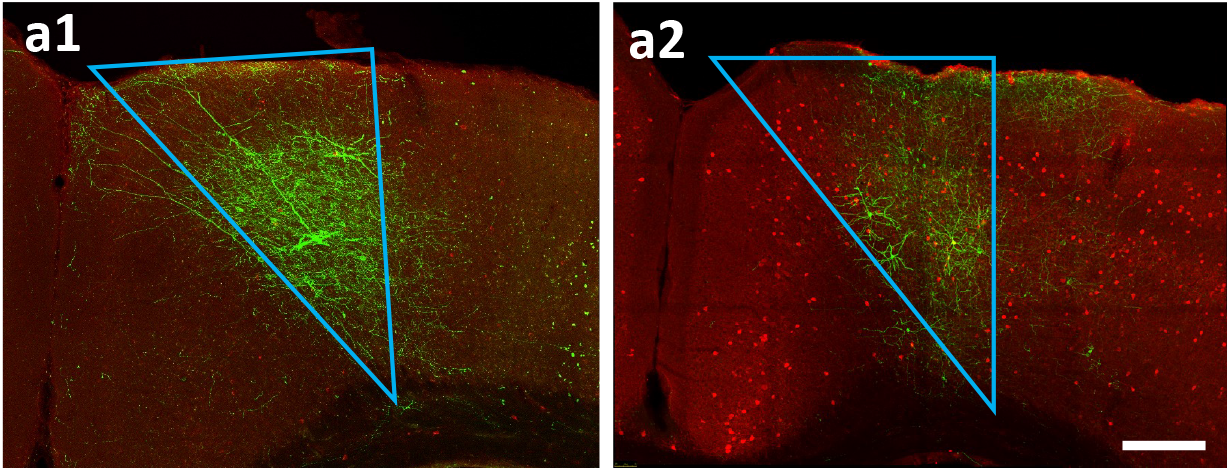
Post-hoc analysis of neurons located in the medial M2. Two examples (A1, A2) of neurons stained with anti-GFP antibody in fixed coronal sections. Parvalbumin staining (red). Triangles represent the medial M2 region. Scale bar 200 μm.

**Supplementary Figure 4:**
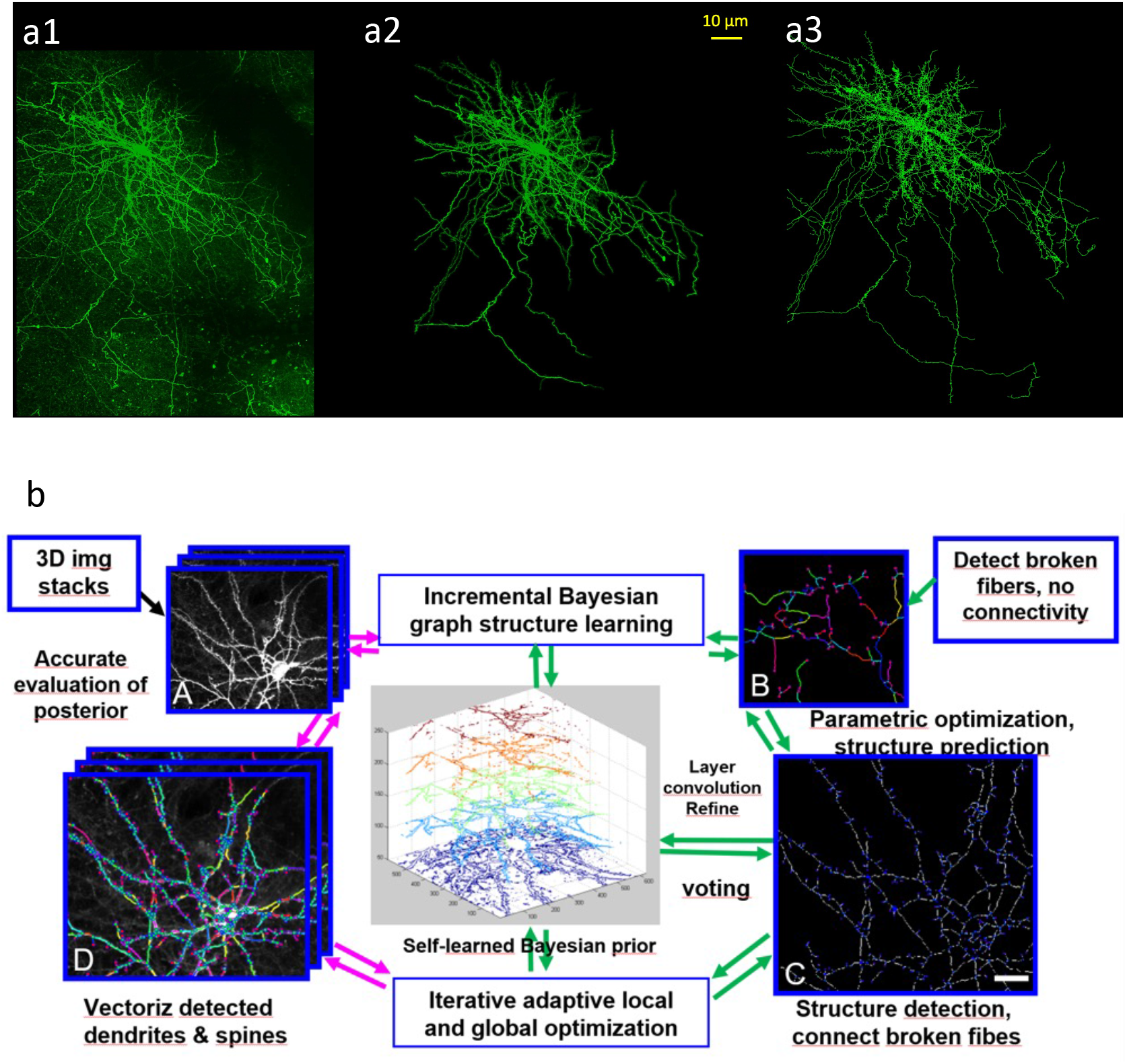
Illustration of image analysis pipeline. (a) Panels showing different steps in image processing. (a1) raw data. (a2) data excised from (a1) using the traced skeleton (a3). (b) Workflow of Bayesian learning-based iterative image analysis strategy. A detailed description can be found in the Supplemental Methods.

**Supplementary Figure 5:**
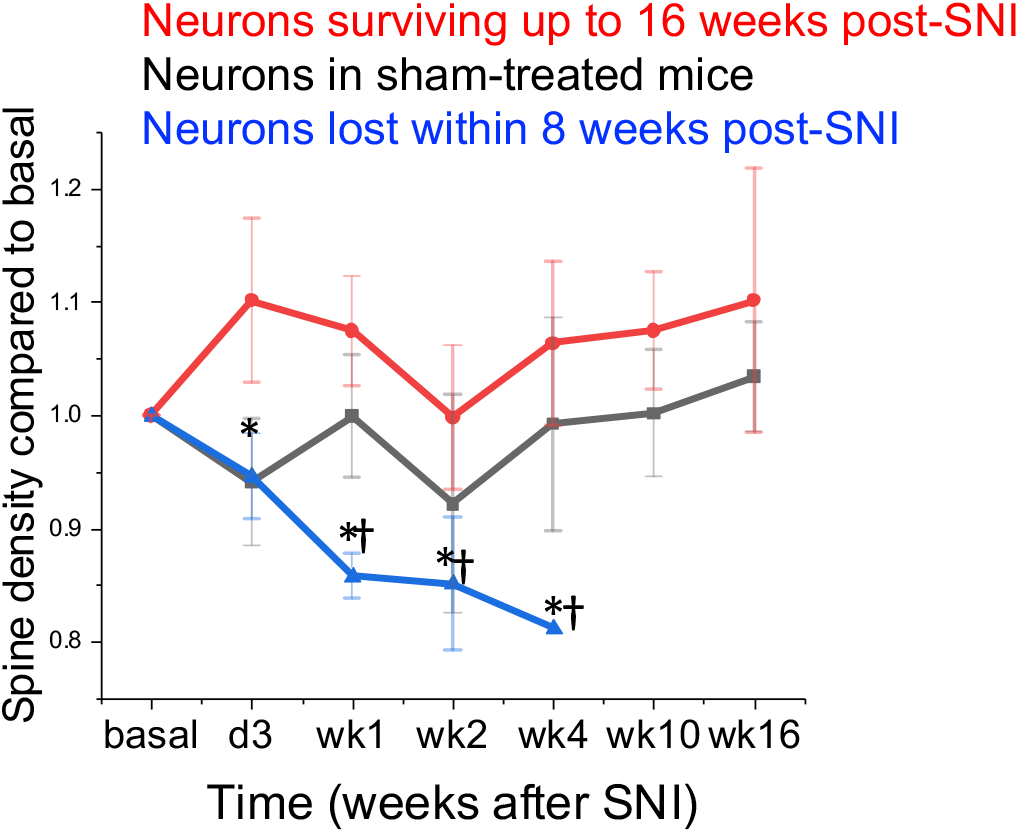
Changes in spine density in neurons prone to die. Comparison of spine density determined for entire neurons over time. Neurons imaged in sham mice (black), neurons imaged in SNI mice surviving 16 weeks, and neurons surviving four weeks or less (blue). Mean ± SEM, N=4 mice in each group, ANOVA with Bonferroni multiple comparison (* = P < 0.05).

