## Supplemental Methods for "Loss of secondary motor cortex neurons in chronic neuropathic pain"

### Supplementary Methods

1. Stitching of individual image stacks into a superstack
2. Automated and adaptive removal of inhomogeneous noisy background
3. Incremental Bayesian learning for accurate neuron 3D tracing and prediction
4. Display of traced cells by means of excised fluorescence
5. Counting of dendritic spines

### 1. Stitching of individual image stacks into a superstack

To image entire neurons in the cortical target volume, we acquired six individual image stacks of approximately 350 µm x 350 µm x 500 µm that were each placed next to each other to form a rectangle (Supplementary Methods Figure 1). Data was acquired at a XY resolution of 85 nm per pixel using a frame size of 4092 x 4092 pixels at 16-bit. For stitching, we reduced the image stacks to half-scale (2046 x 2046 pixels at a pixel size of 170 nm) mainly to limit the size of each 3D stack to become computationally feasible to handle (file size reduction to approximately 18 GB). An overlap of 15-20% was used to allow reliable stitching of the individual stacks into the superstack. Because each stack was acquired in serial order, and due to the long acquisition time required per stack, offsets in the z direction were often encountered that had to be corrected. The resulting stitched image stack typically consisted of 3800 x 5900 x 500 pixels (corresponding to a file size of approximately 100 GB). To achieve this, we used the Fiji 3D stitching plugin ([Image Stitching (imagej.net)]) in combination with a custom-written script.

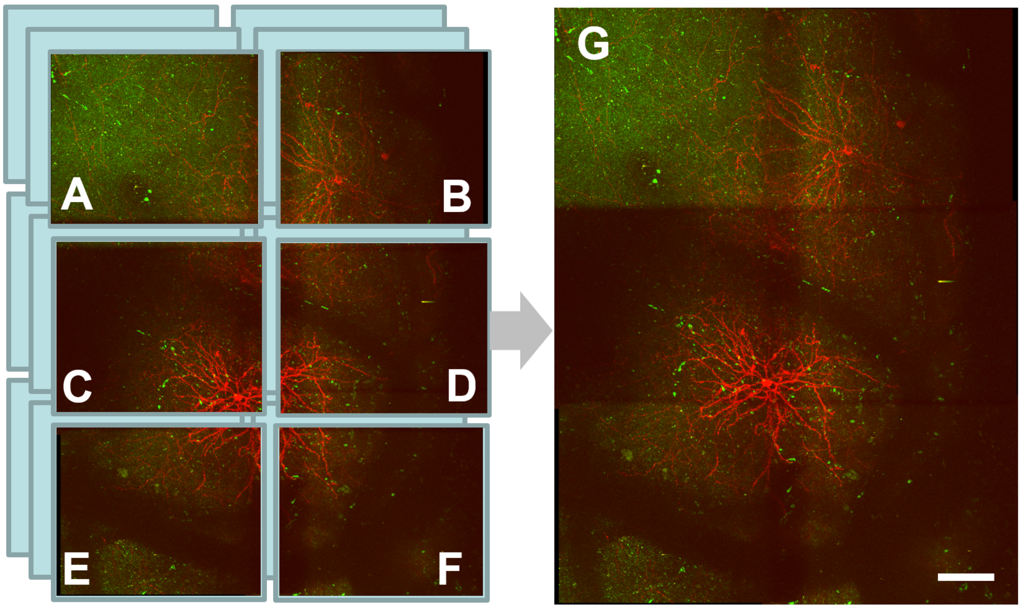


**Supplementary Methods Figure 1 | Automated 3D seamless stitching of six stacks into a superstack. A-F** denote six imaging stacks and their arrangement, overlap region around 15-20%. First A and B are stitched by searching the Z-depth correspondence in 3D space, then correspondence for C and D as well as E and F are established. While searching of correspondence information for pairs of 3D image stacks, the features of dendrites are often crucial for automated correspondence detection. Then seamless stitching and probability-based fusion of all images are performed in an optimization approach for entire 3D image stacks, e.g., least squares methods and its extended methods. **G,** maximum intensity projection (MIP) of a stitched superstack. Scale bar, 100 µm.

### 2. Automated and adaptive removal of inhomogeneous noisy background

Our aim is to automatically detect and segment neuronal processes in 2P image stacks, focusing on dendrites and their spines. While the raw image quality provides a good basis for automated analyses, image processing is required to remove autofluorescence signals. This is particularly relevant for regions at higher depths due to a reduced signal to noise ratio. To achieve this, we developed a custom-written Matlab code that performs automated adaptive background removal. All acquired image stacks were processed as described here and then subjected to the subsequent analysis. The application of this approach is illustrated in Supplementary Methods Figure 2 and detailed in the following paragraph.

Notations and corresponding descriptions

An image is formulated as a weighted undirected graph $G = (V, E, A)$ with nodes $v \in V$, and edges $e \in E \subseteq V \times V$. Each node $v_i$ represents an image pixel $x_i$. An edge $e_{ij}$ connects two nodes $v_i$ and $v_j$ in 8-connected neighborhood system. A is the associated affinity matrix. $A$ is symmetric and non-negative. The diagonal matrix $D = diag(D_{11} \dots, D_{nn})$ is called the degree matrix of graph $G$, where $D_{ii} = \sum_{j=1}^{N} A_{ij}$. To remove two-photon autofluorescence and background, one input image $I$ from an input 3D stack including a foreground image $F$ and a background image $B$. The intensity of the $i$th pixel is assumed to be a linear combination of the corresponding foreground and background intensity value,

$$I_i = \alpha_i F_i + (1 - \alpha_i) B_i, \quad (1)$$

where $\alpha_i$ is the pixel's foreground opacity. In this equation, all quantities on the right side of the compositing equation are unknown and some assumptions on the nature of $F$, $B$ and/or $\alpha$ are needed. To enhance the dendrite tracing performance, we extended an adaptive thresholding and automatic foreground detection algorithm (Zheng, 2006; Bradley, 2007; Wang 2018) to remove the inhomogeneous and noisy background. In this background detection and removal workflow, adaptive image threshold using local first-order statistics is directly computed in a locally adaptive threshold for 2-D grayscale image or 3-D grayscale volume in 16bit 3D image stacks. In Matlab, a function named adaptthresh can be applied. The foreground neural fibers are largely preserved in each local region and the connectivity of neural fibers are also enhanced by means of local adaptive intensity adjustment and enhancement.

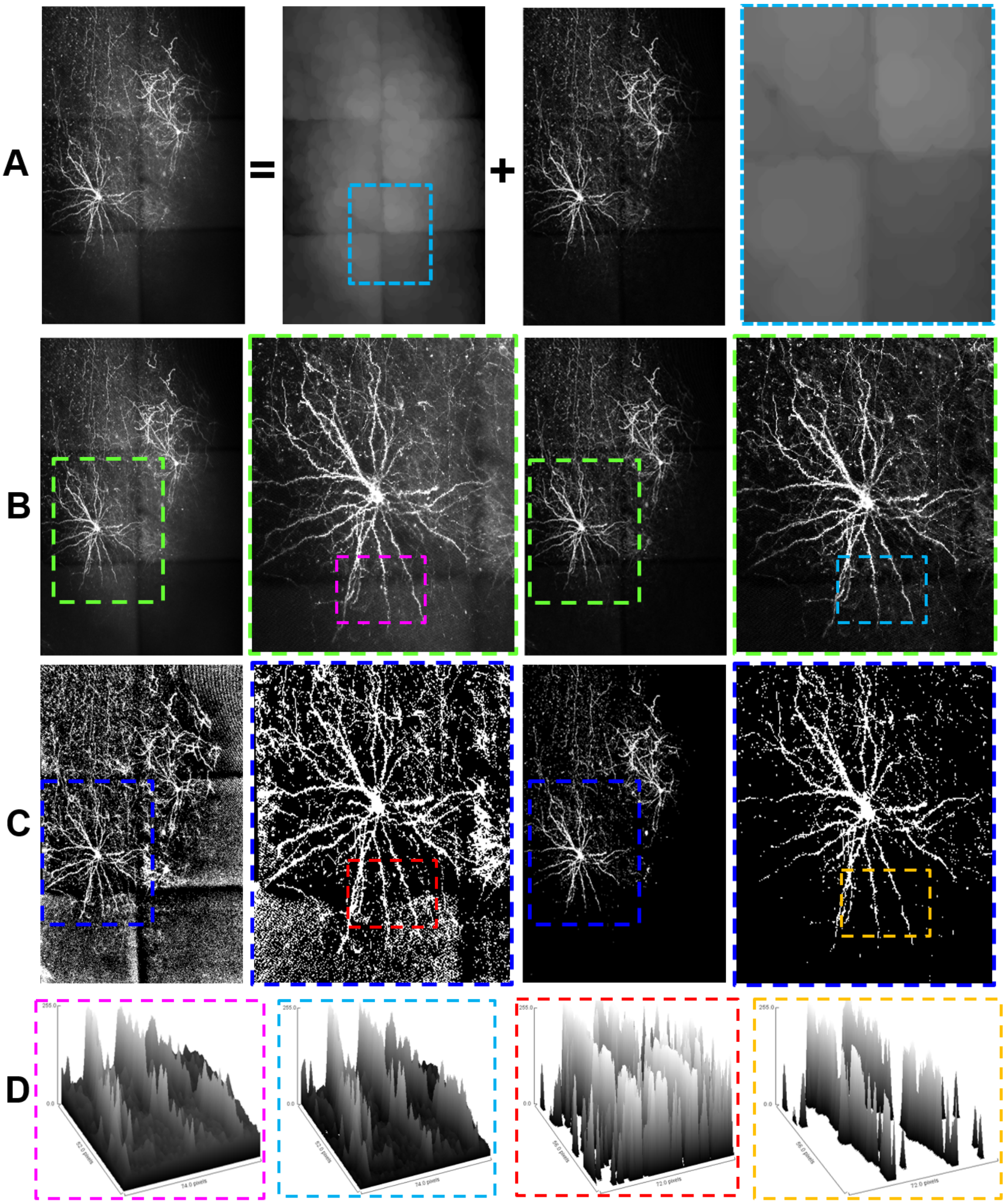


**Supplementary Methods Figure 2 | Adaptive foreground dendrite detection and removal of inhomogeneous background in 3D image stacks. A,** raw image including the labeled neurons and background signals, the latter arise mostly from deep scanning frames. Cyan rectangle delineates area shown enlarged in the right panel. **B,** two left panels show raw image data, right two panel show processed data after removing background. Green rectangle indicates the enlarged view shown in the adjacent panel. Magenta and blue rectangles delineate region of analysis shown in D. **C,** panels shown in B after thresholding the images with the same parameters. Blue rectangle indicates the enlarged view shown in the adjacent panel. Red and orange rectangles delineate region of analysis shown in D. **D.** 3D surface visualization showing the differences in intensities between raw and enhanced images. The enhanced signal to noise ratio of the processed image is much better suited for the subsequent automated tracing algorithm. Scale bar, 50 µm.

#### 3. Incremental Bayesian learning for accurate neuron 3D tracing and prediction

Given that we repetitively acquired the same imaging volumes up to eight times over a time period of 16 weeks, and considering that imaging quality may vary between imaging sessions, we aimed at using a method that would consider information from all imaging sessions and combine it to achieve an overall higher fidelity of tracing. To achieve this goal, we developed a novel incremental Bayesian structure learning and neuronal fiber tracing approach using 16-bit stitched superstacks. In this approach, we first make unsupervised-learning based on multiple superstacks acquired in multiple timesteps. The prior probability and distribution prior can be learned and integrated as a general dendritic structure prior in the Bayesian learning and computing approach. Then the detected and identified dendrites including unconnected or broken dendritic branches are re-computed in the incremental Bayesian structure learning approach in 3D space so that each individual soma and its entire dendritic tree including spines can be traced in an optimization approach (Supplementary Figure 3).

The incremental Bayesian learning approach consists of the following main components:

- Novel Bayesian graph structure learning formula including a self-computed structure prior learning term.
- Adaptive local structure detection and connectivity prediction in a global Bayesian optimization method.
- Incrementally and iteratively improve the tracing posterior results.

These components were implemented in Matlab following established procedures (Zheng and Hellwich, 2006) and extended to include incremental structure learning and prior information reflecting neuronal structure. The performance of this approach is illustrated in Supplementary Methods Figure 3.

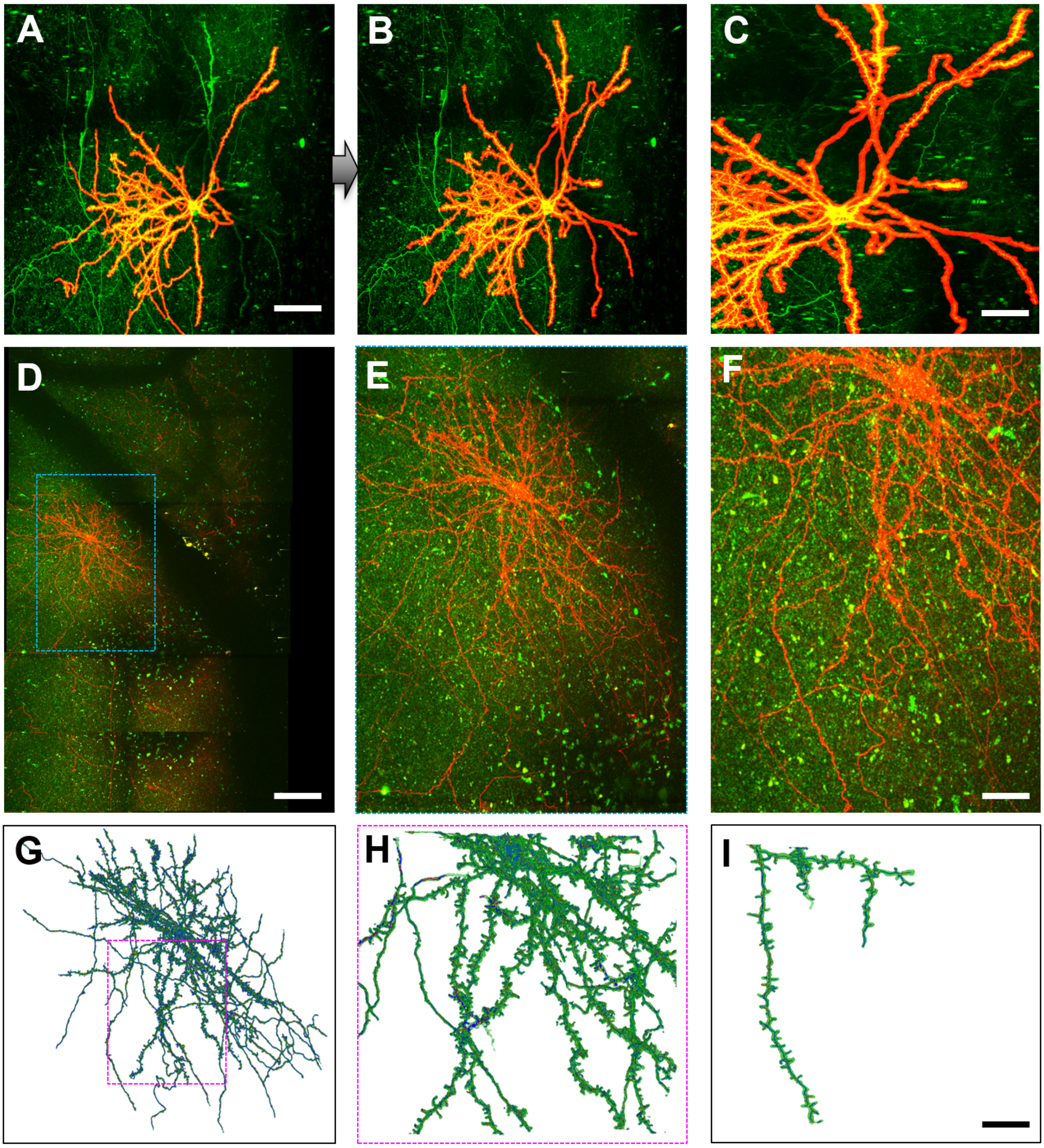


**Supplementary Methods Figure 3 | Illustration of the performance of incremental Bayesian tracing.** **A-C** MIPs showing a traced neuron (red) superimposed on the raw unprocessed data (green). **A** shows the tracing result without using prior information. Note that some faint dendritic structures and entire branches disconnected by a local reduction of the fluorescence signal in the proximal dendrite are not detected. Less intense areas arise from blood vessels positioned in the imaging pathway, thereby obstructing fluorescence signals. **B** shows tracing result of the same cell shown in A, but using Bayesian tracing with prior information. Note that faint dendritic structures and disconnected dendritic branches are now correctly detected. **C**, magnified view of B. **D-E,** MIP showing another neuron traced with our approach (red) superimposed on the raw fluorescence (green). Note less intense, shadow-like areas that arise from blood vessels positioned in the imaging pathway. Blue rectangle denotes area shown enlarged in E, further enlargement in F. **G-H**, Final tracing result of the neuron shown in D-E, illustrating that our approach can achieve highly detailed tracing of the dendritic tree including spines. Shown are MIPs of the excised raw fluorescence generated by using the traced skeleton as a guide to extract the corresponding structures from the raw image (further explanations see paragraph 4 below). Scale bar, 100 µm.

### 4. Display of traced cells by means of excised fluorescence

To achieve the best possible representation of the fully reconstructed neurons and to maintain their original shape and volume information, we used the fully traced neuron represented as a 3D vectorized skeleton obtained from the incremental Bayesian tracing to excise the raw fluorescence signals from the original 16-bit images (Supplementary Figure 4a). This is implemented by storing the width of the traced structure with each point of the single line skeleton. This information can be used to generate a binary volume representation of the neuron for quantitative analysis such as changes in dendritic volume, spine volume, surface, shape and others, or it can be used to excise the raw fluorescence to allow visualization of the neuron without any background noise.

### 5. Counting of dendritic spines

Spine counting was done on the single line skeleton of fully reconstructed neurons. Spines were identified by a junction dot (cyan) connected to a terminal dot (red) as illustrated in Supplementary Methods Figure 4. Neighboring junction dots are not considered to represent spines. Subsequently, all identified spines are counted by the algorithm.

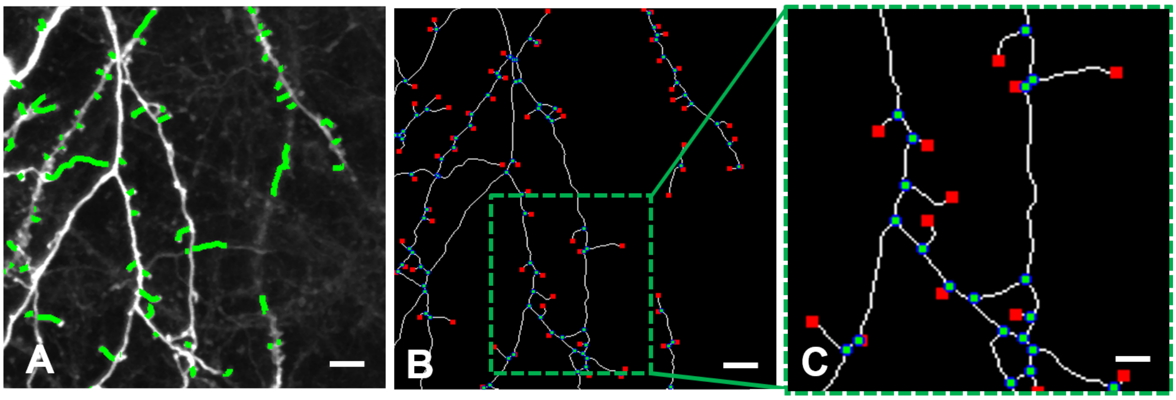


**Supplementary Methods Figure 4 | Illustration of spine detection and counting. A,** Single fluorescence frame superimposed with detected spines (green). **B,** corresponding vectorized skeleton with junction points (cyan) and terminal points (red). Green square denotes area shown enlarged in C. Scale bar, 5 µm.
